# DnaK interacting protein All4144 of *Anabaena* sp. PCC 7120: A thermo-sensor GrpE that acts as a nucleotide exchange factor and binds to DNA

**DOI:** 10.1101/2025.04.14.648684

**Authors:** Sonam Sriwastaw, Ruchi Rai, L.C. Rai

## Abstract

This study investigates the critical role of the dimeric GrpE protein as a thermosensor within bacterial DnaK complexes, with a specific focus on All4144, a GrpE homolog found in *Anabaena* sp. PCC7120. In the present work, we characterized All4144, which possesses both the GrpE domain and a helix-turn-helix (HTH) domain. Our aim was to elucidate the functional significance of the HTH domain by selectively deleting the region encompassing residues 1-67 from the full-length All4144. Our findings reveal that both All4144 and its deletion mutant function as homo-dimeric nucleotide exchange factors for DnaK, enhancing the ATPase and refolding activities of DnaK. Additionally, All4144 plays a role in DNA binding. On the contrary, the deletion mutant lacking the HTH motif exhibits impaired DNA-binding ability, underscoring the importance of the HTH domain. Moreover, All4144 superior cell survival and thermal stability profiles, compared to the deletion mutant. Taken together, our study sheds light on the multifunctional nature of All4144, highlighting its pivotal role in modulating thermal tolerance through interactions with DnaK and DNA.

**Graphical abstract:** 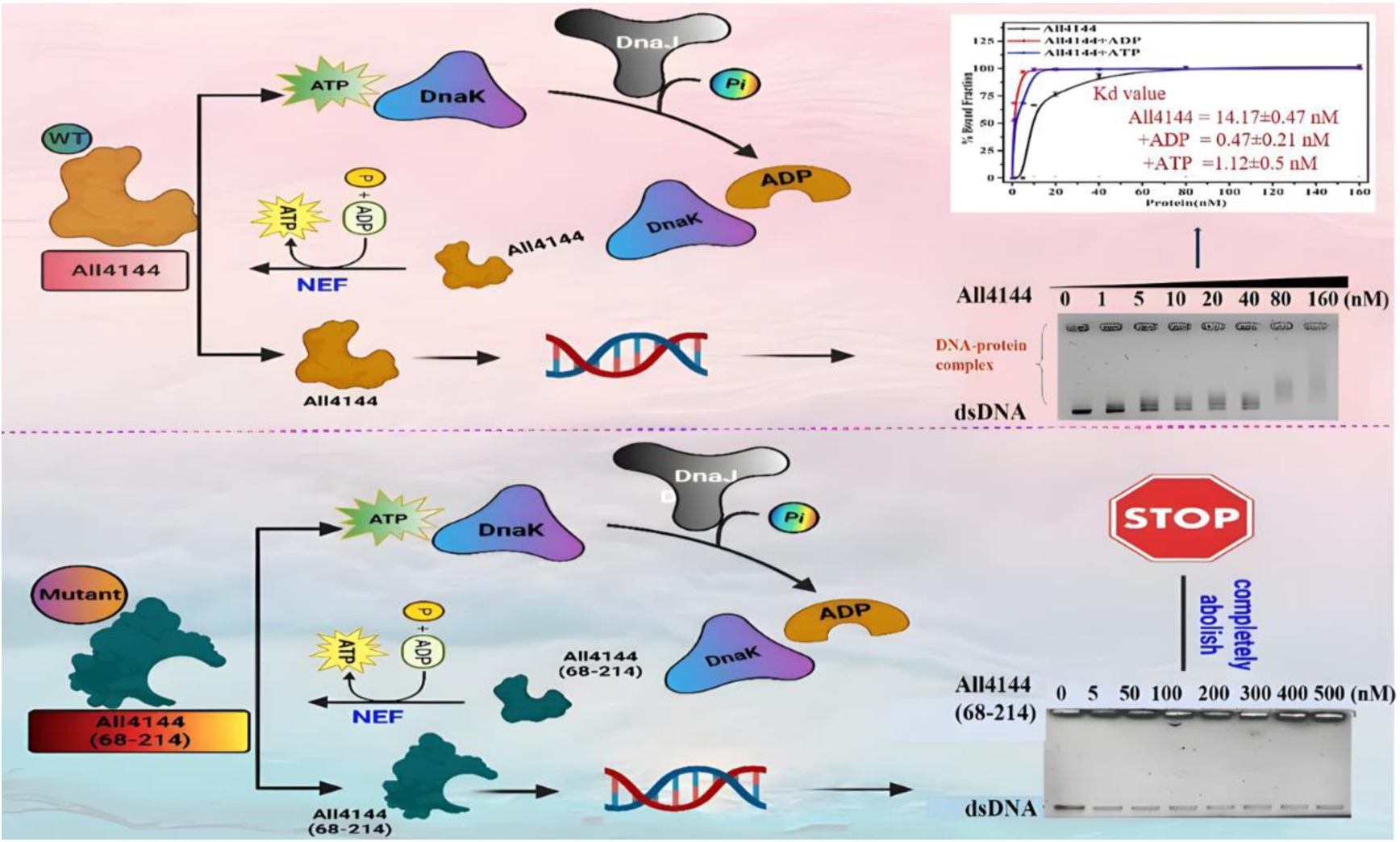

## 1. Introduction

Cyanobacteria are a specialized group of nitrogen-fixing, photosynthetic prokaryotes widely distributed in aquatic and terrestrial habitats (1). Their evolutionary significance relies on their ability to cope with challenging environmental conditions, such as high light intensity, fluctuating temperature, UV-B, and nutrient deprivation (2). Proteomic investigations have unveiled that cyanobacteria respond to such stresses by upregulating the expression of various stress-responsive proteins, particularly heat shock proteins (Hsps) (2). Heat shock proteins, acting as molecular chaperones, play a crucial role in maintaining protein homeostasis by mediating conformational changes and preventing protein aggregation and degradation under abiotic stresses (3). The DnaK (bacterial Hsp70) system is the most studied molecular chaperone, ubiquitously found across all domains of life, and its activity is dependent on two co-chaperones: DnaJ and GrpE (4). Co-occurrence analysis in *Nostoc punctiforme* has revealed the conserved presence of the DnaK-DnaJ-GrpE chaperone system in cyanobacteria (5). DnaJ, an Hsp40 homolog, stimulates ATP hydrolysis and targets substrate protein to ADP-bound DnaK/Hsp70 (6), while GrpE acts as a nucleotide exchange factor (NEF), facilitating ADP/ATP exchange and influencing the longevity of the DnaK-substrate complex (7).

The expression of GrpE in bacteria is a responsive measure triggered by various stressors, including heat shock, nutrient deprivation at low temperatures, and oxidative stresses, underscoring its pivotal role in the intricate web of stress responses (8). This responsiveness emphasizes GrpE’s adaptive nature, aligning with the dynamic needs of the cellular environment under challenging conditions.

In contrast to prokaryotes, eukaryotic homologs of GrpE exhibit noteworthy divergence in sequence identity. Instances include the identification of Mgelp in yeast (9) and GrpEL1 and GrpEL2 in mammalian mitochondria (10). Notably, BAG1 emerges as a functional homolog of GrpE specifically for cytosolic Hsp70s in mammals, further highlighting the versatility and conservation of this essential chaperone system across evolutionary boundaries (11). Structurally, GrpE proteins comprised of three domains: a long paired α-helix, a four-helix bundle-forming domain, and a C-terminus β-sheet domain (12). The long paired α-helices act as a thermosensing domain, undergoing helix-to-coil transitions to prevent thermal denaturation of substrate proteins during heat shock (13). Further, crystal structure analysis of GrpE from *Thermus thermophilus* HB8 revealed that thermosensing action of the GrpE is related to its structural characteristics (14) and mediated by different protein domains i.e. either N-terminus or C-terminus. Despite the well-characterized thermosensing action and function of GrpE as a nucleotide exchange factor, there is no concrete evidence of the diversified biological function of GrpE in prokaryotes. Only fragmentary information is available regarding cyanobacteria e.g. upregulation of a heat shock GrpE protein (Sll0057) in *Synechocystis* sp. when exposed to hexane (8).

Furthermore, GrpE protein has been characterized as a thermosensor in the thermophilic cyanobacterium *Thermosynechococcus elongatus* BP1 and mesophilic *Synechocystis* sp. PCC 6803 (12). Recently, we have observed that DnaJ III of *Anabaena* sp. PCC7120 imparts heat stress tolerance in *E. coli* and stimulates DnaK’s ATPase activity. However, it was not established how activity of the DnaK-DnaJIII chaperone/co-chaperone complex is associated with the co-chaperone GrpE in *Anabaena* sp. PCC7120 (15). This study investigates the functional characteristics of the GrpE homolog, All4144, in *Anabaena* sp. PCC7120. Notably, All4144 exhibits a helix-turn-helix (HTH) domain alongside the conventional GrpE domain. To elucidate the specific function of the HTH domain within GrpE and its implication in DNA binding, we deliberately generated a deletion mutant, All4144(68–214), which lacks the HTH domain. Through a comparative analysis of the full-length All4144 and the deletion mutant using biophysical and biochemical assays, we aim to assess the role of the intact additional HTH domain in conferring multi-functionality, such as thermal tolerance, co-chaperone activity, and DNA binding capability.

## Material and methods

### Cyanobacterial and bacterial strains and plasmids

*Anabaena sp. PCC7120* (hereafter *Anabaena sp.*) was cultivated photoautotrophically in BG-11 medium at 24 ± 2°C, pH 7.4, under daylight fluorescent illumination providing 72 μmol m⁻² s⁻¹ of photosynthetically active radiation (PAR) with a 14 h light/10 h dark cycle. The cultures were stirred three times per day. For cloning and protein overexpression, *Escherichia coli* strains DH5α and BL21(DE3) (Novagen) were used as hosts. Recombinant *E. coli* cultures were maintained in LB medium supplemented with 50 μg/mL kanamycin (16). The pET-28a(+) plasmid (Novagen) was employed for cloning and expression of the target proteins.

### In-silico analysis

Functional domains of All4144 were analyzed using InterPro-Scan (http://www.ebi.ac.uk/interpro) (17) and the Conserved Domain Database (CDD) (http://www.ncbi.nlm.nih.gov) (18). Homology searches were performed using BLASTp from the NCBI database. Multiple sequence alignments were generated using PROMALS-3D (prodata.swmed.edu/promals3d/) (19). The I-TASSER server (20) was employed to generate a three-dimensional homology model of All4144.

### Model Validation and docking

To validate the structural integrity of the generated models, we utilized **PROTEK** and **RAMPAGE** as quality assessment tools (21). **PROTEK** was employed to evaluate the overall structural geometry, ensuring that the model maintained proper folding and stability. Additionally, **RAMPAGE** was used to generate a **Ramachandran plot**, which classified the backbone dihedral angles (ϕ and ψ) of amino acid residues into **favored**, **allowed**, and **outlier** regions. The majority of the residues were located in the favored and allowed regions, confirming that the stereochemistry of the model was within acceptable ranges for high-quality protein structures. Further we have used H-DOCK for protein-receptor docking which uses surface and electrostatic scoring function for protein-protein interactions (22). Best docked pose was selected using the in-built scoring function of H-DOCK.

### Transcript analysis of all4144

Total RNA was isolated from a 100 mL culture of *Anabaena sp.* (OD750 = 0.6) after exposure to arsenic (40 mM, 24 h) (23), heat (42°C, 1 h) (24), NaCl (100 mM, 24 h) (25), UV-B (12.9 mW/cm², 30 min) (26), and cadmium (10 µM, 24 h) (27). cDNA was synthesized from the isolated RNA using the iScript cDNA Synthesis Kit (BioRad). Gene-specific primers were designed using Primer3 (version 4.0, https://primer3.org). Quantitative real-time PCR (qRT-PCR) was performed using 12 ng of cDNA from each sample in triplicate reactions, on a 96-well plate with a total volume of 20 µL, including 10 pmol of forward and reverse primers, and 1× SsoFast EvaGreen qPCR supermix (BioRad). Relative expression of the *all4144* gene was calculated using the ΔCq method (28), normalized to *rnpB* transcript levels. Specificity of the PCR products was confirmed by melting curve analysis.

### Rationale and design of the deletion mutant of All4144

Conserved domain analysis (Fig.1a) identified All4144 as a member of the GrpE family, containing a helix-turn-helix (HTH) domain at the N-terminus (residues 1–67) and a GrpE domain at the C-terminus (residues 68–214). To evaluate the potential role of the HTH domain in GrpE function, an N-terminal deletion mutant, All4144(68–214), was generated by removing amino acids 1–67, as shown in (Fig. 1b-c).

**Figure 1.**
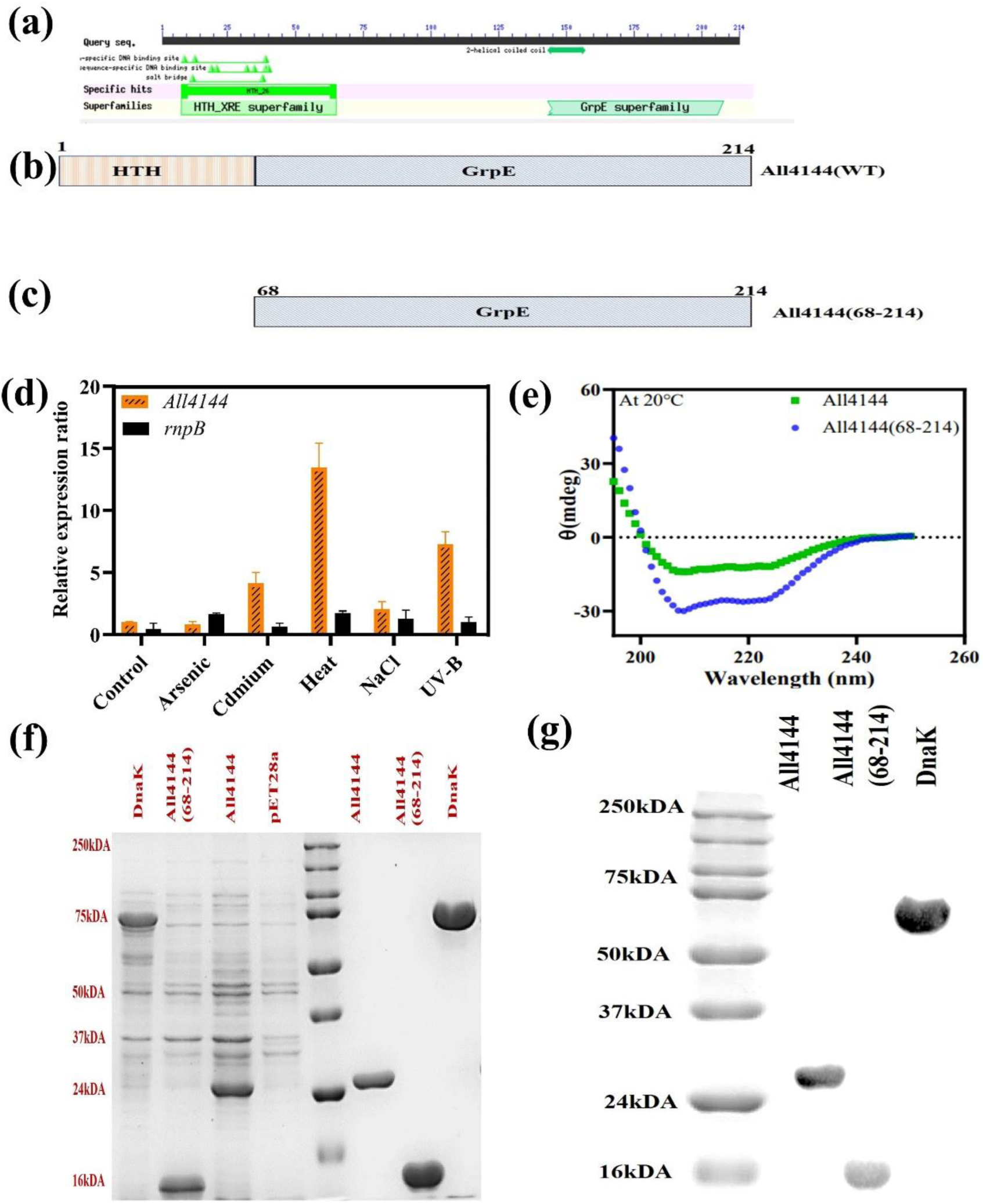
**(a)** Conserved domain analysis of All4144 by Conserved domain database (CDD). (**b)** The full-length protein All4144. **(c)** The N-terminal deletion mutant All4144(68–214). **(d)** Transcript analysis of all4144 in Anabaena sp. PCC 7120 under various abiotic stresses. rnpB gene was used as an internal control for normalizing the variations in cDNA amounts. Relative expression ratio represents the expression of gene of interest in different conditions normalized with respective expressions of the housekeeping gene. **(e)** CD spectra analysis of All4144 and All4144(68–214) at 20 °C. **(f)** Expression analysis of All4144, All4144(68–214) and DnaK by SDS-PAGE. **(g)** Immunoblot detection of purified protein All4144, All4144(68–214) and DnaK. All data are tested in triplicate and the bar indicates standard error (±SE).

### Cloning, expression and purification of All4144 and All4144(68–214)

Genomic DNA from *Anabaena sp.* was isolated and used for PCR amplification of *all4144 and all4144*(68–214), which were cloned into the pET28a(+) plasmid at the NdeI and XhoI restriction sites. The recombinant plasmids were transformed into *E. coli* BL21, and protein expression was induced with 0.2 mM IPTG. Recombinant cells carrying *all4144* and *all4144*(68–214) were grown in LB broth containing 25 µg/mL kanamycin at 37°C with shaking at 160 rpm until the optical density at 600 nm reached 0.3. Protein expression was induced by adding 0.2 mM IPTG, followed by growth at 18°C for 14 h at 160 rpm. Cells were harvested by centrifugation at 15,000×g for 15 min at 4°C, and resuspended in 8 mL of cold lysis buffer (20 mM Tris-HCl, 500 mM NaCl, 40 mM imidazole, pH 7.8). Cell lysis was performed using ultrasonic treatment, and the lysate was clarified by centrifugation at 13,000×g for 50 min at 4°C. His-tagged proteins were purified from the supernatant using a His-Trap Nickel-NTA column (Bio-Rad). Eluted protein fractions were analyzed by SDS-PAGE (12%), and fractions with >95% purity were pooled and concentrated using a 10 kDa Amicon concentrator (Millipore). Protein concentrations were determined by absorbance at 280 nm.

### Cell survival study

The impact of heat stress was assessed using a spot assay, as described previously (29). In brief, recombinant cells and empty vector control (pET28a) were cultured in LB medium containing 50 µg/mL kanamycin at 37°C. Once the cultures reached an OD_600_ of 0.6, they were induced with 0.2 mM isopropyl β-D-1-thiogalactopyranoside (IPTG) and incubated for an additional 5–6 h at 16°C. The IPTG-induced *E. coli* cells were then heat-treated at 50°C for 30, 60, and 90 min. Cultures were serially diluted (10⁻¹, 10⁻³, and 10⁻⁵), and 5 µL of each dilution was spotted onto LB-agar plates containing kanamycin. The spot assay results were further validated by SDS-PAGE analysis, where cells grown at 37°C and 50°C were harvested, proteins were isolated, and the expression was evaluated by SDS-PAGE. To ensure accurate comparisons, we used the housekeeping gene β-actin as a reference control.

Quantitative analysis of the impact of heat stress on genetically modified *E. coli* BL21 (DE3) strains was conducted using the colony-forming unit (CFU) count method. Following the same growth and induction protocol as the spot assay, a 50 μl aliquot of a culture diluted to 10^−3^ was evenly spread onto LB-kanamycin plates. Bacterial colonies were counted, and CFU was calculated using the formula: CFU = (number of colonies × dilution factor) / volume of culture plated.

### Intracellular reactive oxygen species (ROS) measurement

Intracellular reactive oxygen species (ROS) generation under heat stress conditions was measured using 2,4-dichlorofluorescein diacetate (H₂DCFDA) following the protocol by Jakubowski and Bartosz (30). Overnight cultures of *E. coli* BL21 (DE3) cells harboring either an empty vector (pET-28a) or recombinant plasmids, pET-28a-all4144 and pET-28a-all4144(68–214), were grown in LB medium containing 50 µg/mL kanamycin. A 100 µL aliquot of these cultures, at OD600 = 0.6, was inoculated into 50 mL of LB medium supplemented with 0.2 mM IPTG. Cultures were subjected to heat stress at 50°C or maintained at 37°C as a control, shaking at 200×g for 60 min. After incubation, cells were harvested by centrifugation, washed, and resuspended in PBS buffer (pH 7.4). ROS generation was assessed by adding 10 µM H₂DCFDA (dissolved in ethanol) to the cell suspension, followed by incubation at 37°C for 30 min. ROS production was quantified using a Cary Eclipse LS-5B spectrofluorimeter with excitation at 490 nm and emission at 520 nm.

### Circular dichroism (CD) and DSC analysis

Circular dichroism (CD) spectra for full-length All4144 and its deletion mutant, All4144(68–214), were recorded in the far-UV region (190–250 nm) at a concentration of 5 µM in 25 mM phosphate buffer (Buffer A). CD measurements were conducted on a Jasco J-810 spectropolarimeter, using a 2-mm path length quartz cell. Temperature was incrementally varied from 20°C to 80°C, with a response time set at 50 nm/s. Prior to each sample measurement, a baseline spectrum of Buffer A was obtained and subtracted. Additional thermal scans were performed at 222 nm to monitor changes in α-helical content. For each sample, three scans were collected and averaged to improve data reliability. Melting temperatures (T_m) were determined from spectral data at specific wavelengths indicative of secondary structural elements and were fit to a sigmoidal curve (Boltzmann distribution), indicating cooperative unfolding behavior consistent with a two-state mechanism, where structural transitions occur concertedly across the molecule.

Differential scanning calorimetry (DSC) was performed according to a previously described method (31), with modifications. In brief, purified proteins in buffer C were analyzed using a DSC instrument (Mettler Toledo 822E, India). Samples (5 µM) were weighed into sealed aluminum crucibles, and thermal analysis was conducted in a nitrogen-depleted atmosphere over a temperature range of 0–200°C at a heating rate of 10°C/min. This method was employed to measure the excess heat capacity as a function of temperature and to calculate the calorimetric enthalpy of thermal unfolding (ΔH_cal) for both proteins (32).

### Cross-linking by glutaraldehyde

Glutaraldehyde crosslinking of proteins in their native state was conducted as previously described (33). Briefly, 5 µM of All4144 and its deletion mutant proteins were diluted in buffer A to a final reaction volume of 25 µL. Protein solutions were incubated at 37°C for 8 min, either in the absence or presence of 1.5 µL freshly prepared 0.025% glutaraldehyde solution. Following incubation, an equal volume of 2X Laemmli sample buffer was added, and the mixtures were heated at 90°C for 3 min. The crosslinked proteins were separated by 12% SDS-PAGE and visualized with Coomassie brilliant blue staining.

### Refolding assay

Malate dehydrogenase (MDH) was chemically denatured by incubating it for 24 h in a buffer containing 6 M guanidine hydrochloride (GdnHCl), 50 mM Tris-HCl (pH 7.5), 5 mM MgCl₂, and 2 mM dithiothreitol (DTT). Refolding of denatured MDH (5 nM) was performed using a previously described method (34). The denatured MDH was diluted in a buffer consisting of 100 mM Tris-HCl (pH 7.5), 3 mM glucose 6-phosphate, 5 µM ATP, and 150 mM NAD. The refolding reaction included DnaK (1 µM), DnaJ (1 µM), and either All4144 or the deletion mutant protein (2 µM), and was incubated for varying durations. Kinetics of MDH refolding were monitored using a spectrophotometer at a wavelength of 340 nm.

### ATPase activity assay

DnaK ATPase activity was measured with modifications based on the method described by Lanzetta et al. (35). Briefly, a reaction mixture containing 1 µM recombinant DnaK, 1 µM DnaJ, and 2 µM of either full-length All4144 or All4144(68–214) proteins was incubated in ATPase assay buffer C, which comprised 30 mM Tris, 50 mM KCl, 0.6 mM dithiothreitol (DTT), 2 mM MgCl₂, 1 mM ATP, and 10% (v/v) glycerol. A control reaction was established that excluded DnaJ, All4144, and the deletion mutant proteins, including only DnaK in the buffer. Reactions were initiated by the addition of 1 mM ATP and incubated at 25 °C. At various time intervals, reactions were quenched by adding 4 M formic acid. Values were monitored using a spectrophotometer at a wavelength of 650 nm.

### Tryptophan quenching analysis DnaK

The intrinsic tryptophan fluorescence of DnaK was measured using a Perkin-Elmer LS-5B Cary Eclipse spectrophotometer. Nucleotide-free DnaK, at a concentration of 1 µM, was analyzed at 25 °C in a buffer composed of 50 µM Tris-HCl (pH 7.5), 100 mM KCl, 5 µM MgCl₂, 2 mM EDTA, and 2 mM dithiothreitol (DTT). Emission spectra were recorded with a fixed excitation wavelength of 290 nm. For the experiments involving ADP, ATP, All4144, and the deletion mutant protein, 2 µM of each component was added to the DnaK sample. ATP or ADP was introduced from a stock solution (100 µM), and the mixture was incubated for 5 min in the cuvette at room temperature. Emission spectra were then recorded over a wavelength range from 300 nm to 400 nm (36).

### Electrophoretic mobility shift assay

Electrophoretic mobility shifts assays (EMSAs) were conducted with minor modifications based on previously described methods (37). Briefly, 1 µM protein samples were incubated with double-stranded DNA (dsDNA) (501 bp) and single-stranded DNA (ssDNA) (60 bp) in a total volume of 12 µL of binding buffer (50 mM Tris, pH 7.45; 750 mM KCl; and 0.5 mM dithiothreitol [DTT]), in the absence and presence of 1 mM ATP and ADP. The samples were incubated for 15 min at 25 °C before being loaded onto 1% agarose gels for electrophoresis in 1X TBE buffer for 90 min. Following electrophoresis, the gels were stained with ethidium bromide and imaged using a gel documentation system to visualize the DNA-protein complexes.

### Statistical analysis

All experiments were performed in triplicate, and statistical significance was assessed using analysis of variance (ANOVA) with SPSS software (SPSS Inc., Chicago, IL, USA). Graphical representations of the data were created using GraphPad Prism 8 and Origin 9 (Origin Lab, USA), with values expressed as mean ± standard error of the mean (SEM).

## 3. Result

### Transcript analysis of all4144 under multiple abiotic stresses using quantitative RT-PCR

Transcript profiling of *all4144* was conducted under various abiotic stresses, including heat, UV-B, cadmium, arsenic, and salt. The expression of *all4144* was significantly upregulated under heat, cadmium, and UV-B stress, showing increases of approximately 13.4 ± 1.9, 7.2 ± 0.9, and 4.1 ± 0.8 folds, respectively, compared to the control (Fig. 1d). In contrast, the fold changes in response to arsenic and salt stress were only 0.8 ± 0.2 and 2.0 ± 0.59, respectively.

### Purification and Secondary Structure of All4144 and All4144(68–214)

The open reading frames of *all4144* and its deletion mutant *all4144*(*68–214*) were amplified to investigate the products of the heat-inducible gene. Transformants of *E. coli* BL21(DE3) were generated, resulting in substantial production of soluble proteins with molecular weights of 24 kDa and 16 kDa, respectively (Fig. 1f). Western blot analysis using an anti-His antibody confirmed successful protein expression (Fig. 1g). Additionally, Far-UV circular dichroism (CD) analysis indicated that both All4144 and All4144(68–214) exhibit a significant α-helical content and an approximate β-pleated component, confirming that both proteins are well-folded (Fig. 1e).

### Cell survival study

Cell survival analysis of recombinant *E. coli* strains and the control (pET28a) was assessed using a spot assay and colony-forming unit (CFU) count. Spot assays were performed at 37°C and 50°C for 30, 60, and 90 minhou. The results indicated that both recombinant strains, BL21/pET-28a-all4144 and BL21/pET-28a-all4144(68–214), exhibited increased heat resistance compared to the vector control (BL21/pET-28a). Notably, the wild-type *all4144* strain demonstrated superior growth under heat stress conditions compared to the deletion mutant BL21/pET-28a-all4144(68–214) (Fig. 2a-b).

**Figure 2.**
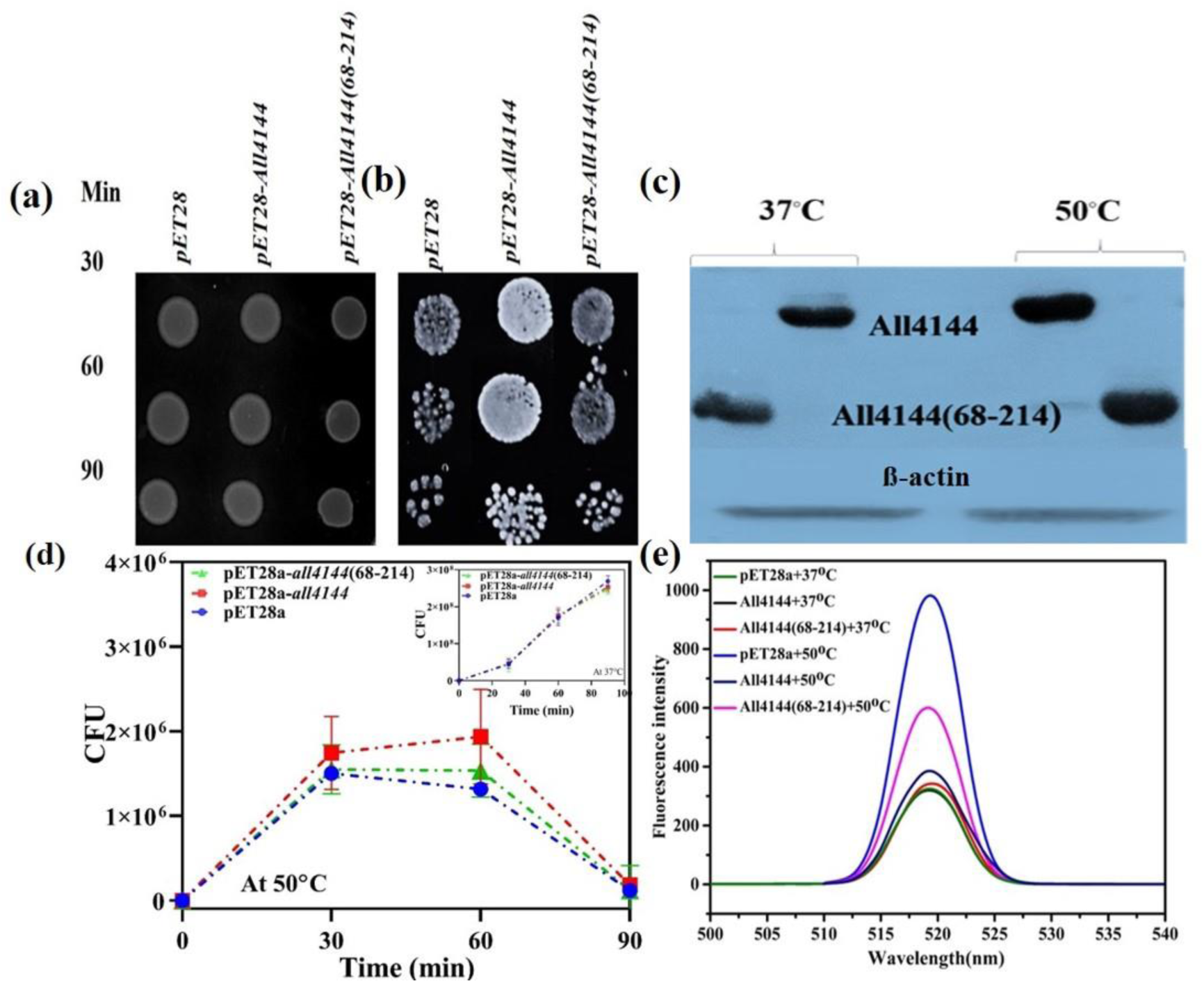
**(a-b)** Measurement of heat stress tolerance using spot assay of E. coli strain BL21 transformed with empty vector (BL21/pET-28a) and recombinant plasmid {BL21/pET-28a-all4144 and all4144(68–214)}. **(c)** Expression analysis of All4144 and All4144(68–214) under cell stress conditions by SDS-PAGE. **(d)** CFU/ml of (BL21/pET-28a) and BL21/pET-28a-all4144 and all4144(68–214)} under heat stress. The data contain the mean of three replicates, and the bars show the standard error (S.E.). **(e)** Reactive oxygen species (ROS) generation analysis of E. coli BL21 cells carrying the control vector (BL21/pET-28a) and recombinant vector (BL21/pET-28a-all4144 and BL21/pET-28a-all4144(68–214) under heat stress using 2,4 dichlorofluorescein diacetate (H_2_DCFDA).

Quantitative validation of these observations was obtained through CFU count analysis (Fig. 2d). CFU results showed comparable spot densities for BL21/pET-28a and the recombinant cells at 37°C. However, at 50°C, BL21/pET-28a-all4144 exhibited a substantial increase in spot density relative to the control, with fold changes of approximately 1.4, 2.8, and 5 at 30, 60, and 90 min, respectively. In contrast, BL21/pET-28a-all4144(68–214) cells showed fold changes of 1, 1.6, and 1.2, indicating a less pronounced response to heat stress than BL21/pET-28a-all4144.

SDS-PAGE analysis of protein expression under heat stress revealed higher protein expression levels at 50°C compared to 37°C and the control (Fig. 2c), indicating that both proteins (All4144 and the deletion mutant) upregulate in response to heat stress.

To further evaluate the reactive oxygen species (ROS) generation following the spot assay and CFU count. We found that both BL21/pET-28a-all4144 and BL21/pET-28a-all4144(68–214) exhibited decreased H_2_DCFDA dye intensity compared to the control (BL21/pET-28a), indicating reduced ROS production (Fig. 2e). However, BL21/pET-28a-all4144 demonstrated better heat tolerance than the deletion mutant, suggesting that the HTH domain impacts on cell survival under heat stress conditions.

### Thermal stability of All4144 and All4144(68–214)

The thermal stability of the full-length protein All4144 and its deletion mutant, All4144(68–214), was examined using circular dichroism (CD) spectroscopy and differential scanning calorimetry (DSC). CD spectroscopy revealed that both proteins maintained their secondary structure up to approximately 60 °C, with All4144 showing a higher α-helical content compared to the deletion mutant (Fig. 3a-b). The thermal unfolding profile was tracked by the decline in ellipticity at 222 nm, a wavelength sensitive to helical content, with All4144 exhibiting a main thermal transition at ∼65.24 °C, whereas All4144(68–214) displayed a lower thermal transition midpoint at ∼60.78 °C. Additionally, an early thermal transition observed between 30-40 °C in both proteins may indicate a potential physiological role at this temperature range (Fig. 3c-d). DSC analysis provided further insights, revealing two distinct thermal transitions in both proteins. For the full-length All4144, transitions were observed at 37.50 ± 0.07 °C and 101.17 ± 0.08 °C, suggesting significant structural stability across a wide temperature range. In contrast, All4144(68–214) displayed transitions at 39.91 ± 0.03 °C and 76.30 ± 0.05 °C, indicating a reduced thermal stability in the deletion mutant. Together, these findings from CD and DSC analyses suggest that All4144 has enhanced thermal stability compared to All4144(68–214), with implications for its structural integrity and function under varying thermal conditions (Fig. 3e-f).

**Figure 3.**
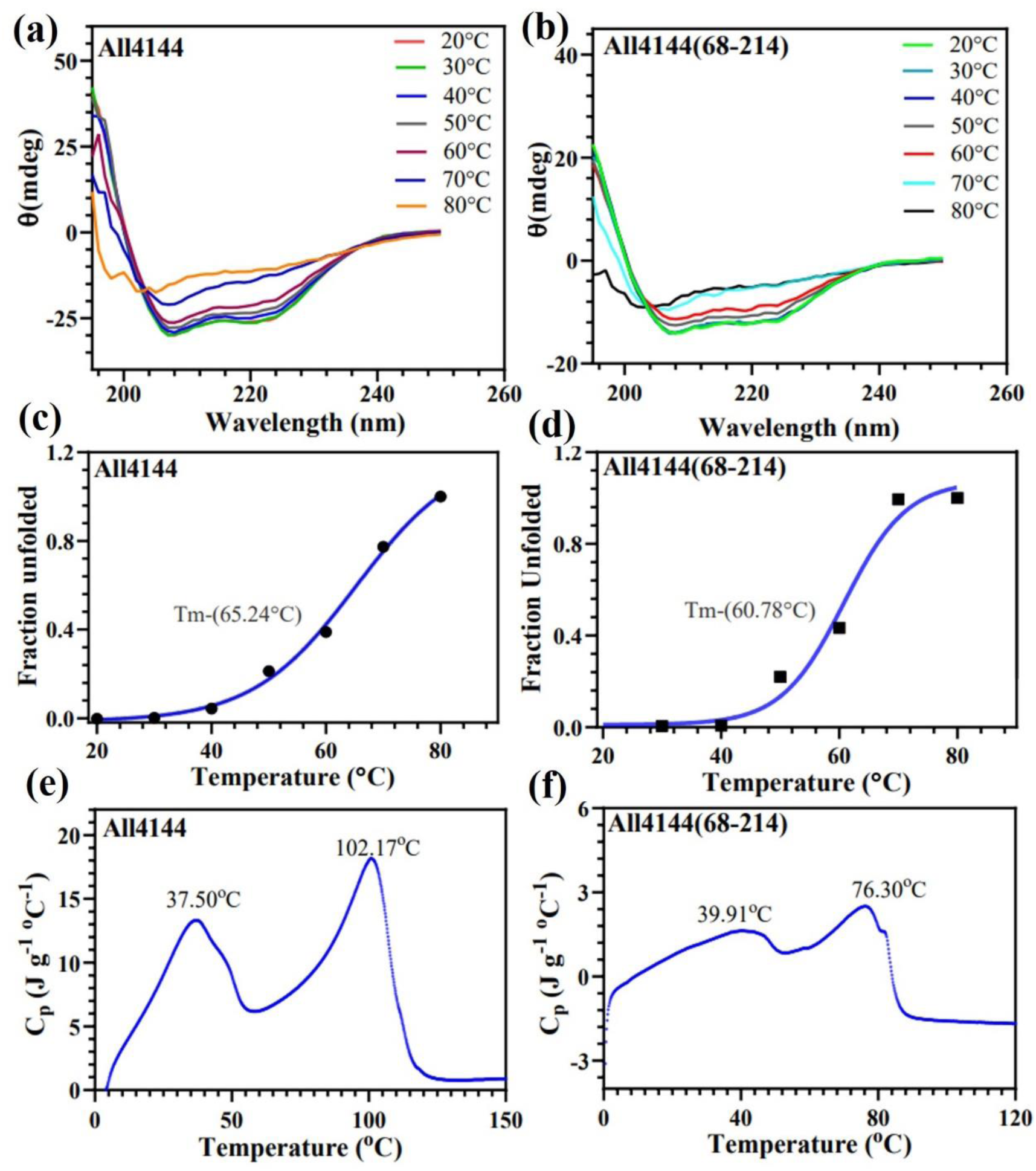
Comparative Analysis of Secondary Structure and Thermal Stability of All4144 and All4144(68–214). **(a-b)** The secondary structure and thermal stability of the full-length protein All4144 and its deletion mutant, All4144(68–214), were characterized using circular dichroism (CD) spectroscopy. Far-UV CD spectra were recorded between 190–260 nm over a temperature range from 20 °C to 80 °C, allowing for detection of secondary structural elements in both All4144 and All4144(68–214). **(c-d)** CD ellipticity measurements at 222 nm were recorded to monitor the thermal denaturation profiles of each protein. The decrease in ellipticity with increasing temperature reflects changes in helical content, providing insights into structural stability under thermal stress. **(e-f)** Differential scanning calorimetry (DSC) was performed to analyze the thermal unfolding properties of All4144 and All4144(68–214). The DSC thermograms display endothermic peaks corresponding to thermal transitions, with peak temperatures indicating the unfolding points (Tm values) for each protein.

### All4144 and All4144(68–214) are dimeric proteins

To examine the dimerization of purified proteins All4144 and All4144(68–214), were incubated without a cross-linking reagent. Both proteins were identified as monomers, with molecular weights of approximately 24 kDa and 16 kDa, respectively. Following the addition of glutaraldehyde, both All4144 and All4144(68–214) formed homodimers, resulting in molecular weights of ∼50 kDa and ∼32 kDa, respectively (Fig. 4a). These results indicate that the deletion of the HTH domain in All4144(68–214) does not disrupt the dimerization capability of All4144. The consistent formation of stable homodimers in both proteins underscores the importance of dimeric structures in elucidating their functional characteristics.

**Figure 4.**
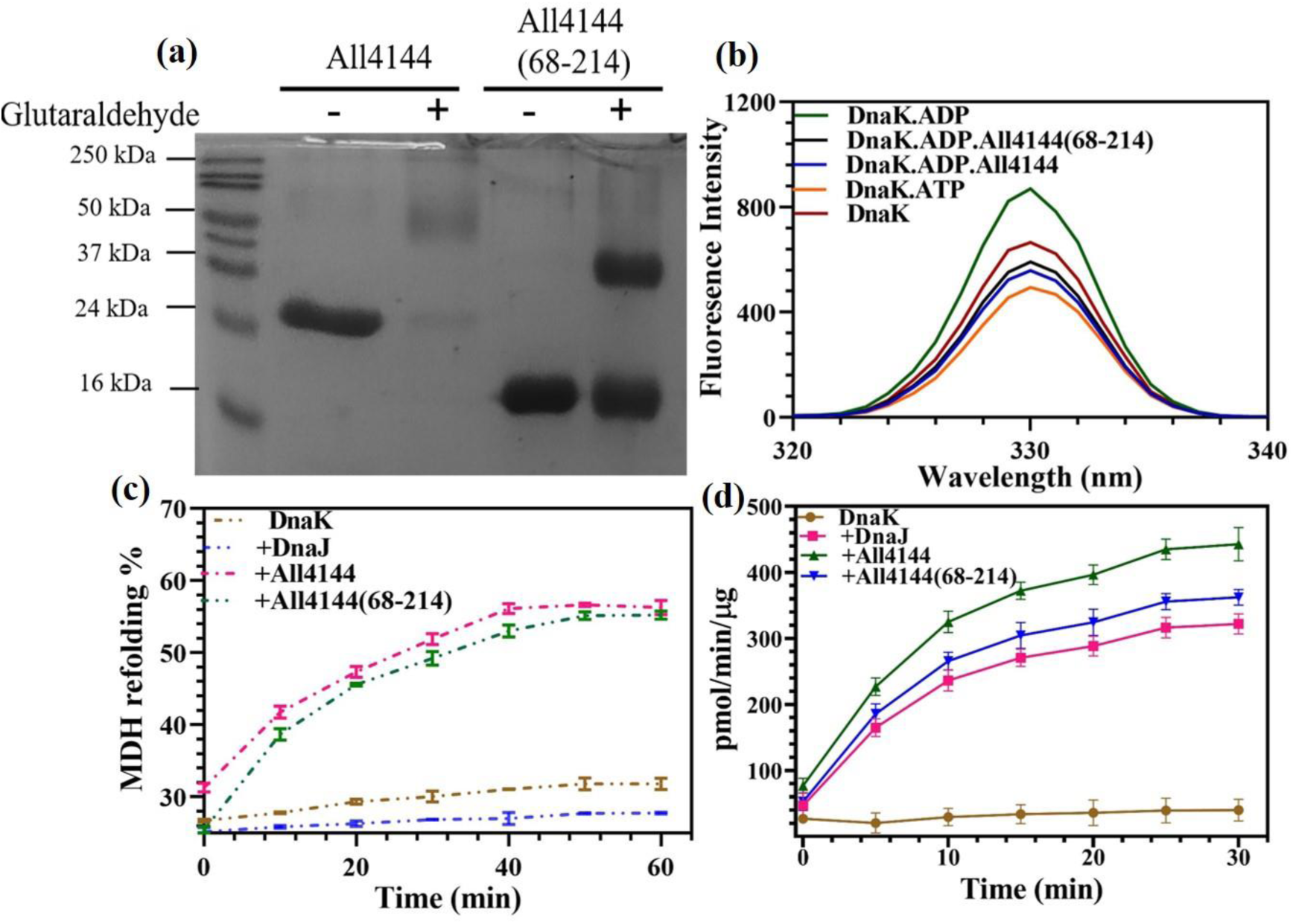
**(a)** Dimerization activity analysis of All4144 and All4144(68–214) protein in solution using glutaraldehyde crosslinking. The data shows the mean of three replicates, and the bars show the standard error (S.E.). **(b)** The fluorescence spectra of ADP. DnaK and ATP. DnaK was compared with that of ADP. DnaK.All4144 and ADP. DnaK. All4144(68–214). DnaK, ATP/ADP, and {All4144 and All4144(68–214)} concentrations were 1 µM, 1 mM, 2 µM and 2 µM, respectively. **(c)** Chemically denatured MDH was refolded in the presence of recombinant DnaK, All4144, All4144(68–214). The ability to refold was measured as a function of enzyme activity by monitoring the reduction of NAD at 340 nM **(d)** The ATPase activity of DnaK in the presence of All4144 and All4144(68–214) was measured at OD_630_ at different time points. The data contain the mean of three replicates, and the bars show the standard error (S.E.).

### All4144 and All4144(68–214) form intermediate structure between ATP-DnaK and ADP-DnaK

The fluorescence emitted by tryptophan residues within DnaK was utilized to assess the impact of All4144 and All4144(68–214) on the structural arrangement of DnaK, both in the presence and absence of ADP/ATP. Our study revealed that ATP caused a reduction in DnaK fluorescence, whereas ADP alone did not produce the same effect. Moreover, the addition of both All4144 and All4144(68–214) to the DnaK.ADP mixture led to a decrease in the intrinsic fluorescence of DnaK. Consequently, based on the intrinsic fluorescence spectra, both All4144 and All4144(68–214) induced an intermediate structural state between DnaK bound to ATP and DnaK bound to ADP. However, All4144 resulted in a greater quenching of DnaK fluorescence spectra compared to its All4144(68–214), indicating its higher affinity for DnaK. These findings extend our comprehension of the molecular interactions and conformational alterations that occur upon the binding of All4144 and its deletion mutant to DnaK (Fig. 4b).

### All4144 and All4144(68–214) function as true co-chaperons

In this study, we investigate the role of All4144 and All4144(68–214) in enhancing the refolding efficiency of denatured malate dehydrogenase (MDH) mediated by DnaK. Our findings reveal the physiological significance of All4144 in modulating DnaK’s chaperone function, particularly concerning protein refolding. In the absence of chaperones, less than 25% of MDH was reactivated, which served as a control. The addition of DnaK alone increased this to only 29%. However, the presence of both All4144(68–214) and All4144 markedly enhanced refolding efficiency, achieving 53.79% and 56.53%, respectively (Fig. 4c). These results indicate that the deletion mutant still significantly enhances the refolding of MDH mediated by DnaK. Notably, the deletion of the N-terminal region does not impair MDH’s refolding activity, allowing it to function effectively as a genuine co-chaperone.

### All4144 and All4144(68–214) stimulate the ATPase activity of DnaK

We assessed the rates of DnaK-catalyzed ATP hydrolysis in the presence or absence of DnaJ, All4144, and All4144(68–214). DnaK exhibited a low hydrolysis rate of 0.009 nMol/µg protein/min ATP hydrolyzed (Fig. 4d). The addition of DnaJ enhanced the rate roughly by 8-fold. Interestingly, when DnaJ was combined with All4144 or All4144(68–214), the rates of ATP hydrolysis by DnaK increased by 11-fold and 9-fold, respectively (Fig. 4d). Furthermore, the fact that the deletion mutant still stimulates the ATPase activity of DnaK indicates that the N-terminal HTH domain does not completely affect its function, leading it to work as a nucleotide exchange factor.

### All4144 is a DNA binding protein

The DNA-binding capabilities of All4144 and its deletion mutant All4144(68–214), were examined using an electrophoretic mobility shift assay (EMSA). The results revealed that full-length All4144 formed nucleoprotein complexes, as indicated by a retardation in DNA mobility in the gel. Notably, full-length All4144 exhibited nonspecific binding with non-specific dsDNA significantly. In the presence of ATP and ADP, DNA demonstrated substantial stimulation compared to the protein alone (Fig. 5a-c). The affinity of All4144 for dsDNA was found to be Kd = 14.17 ± 0.47 nM. Additionally, the binding of All4144 to dsDNA increased nearly 12.6-fold in the presence of ATP (Kd = 1.12 ± 0.05 nM) and 31-fold in the presence of ADP (Kd = 0.45 ± 0.21 nM) (Fig. 5e). These findings suggest that All4144 non-specifically binds with dsDNA. Additionally, All4144 did not bind to ssDNA, even at higher concentrations of the protein used (Fig. 5f-h), confirming a preference for dsDNA over ssDNA in both the presence and absence of ADP/ATP. In contrast, the deletion mutant All4144(68–214) exhibited a complete loss of binding ability to both dsDNA and ssDNA (Fig. 5d). This was evident from the inability of All4144(68–214) to form nucleoprotein complexes with dsDNA, even at high protein concentrations up to 500 nM (Fig. 5d). These results demonstrate that the helix-turn-helix (HTH) domain at the N-terminus of All4144 is essential for binding to dsDNA. The removal of the HTH domain leads to the complete loss of the DNA-binding ability of the protein, emphasizing the critical role of the HTH domain in mediating the interaction between All4144 and dsDNA.

**Figure 5.**
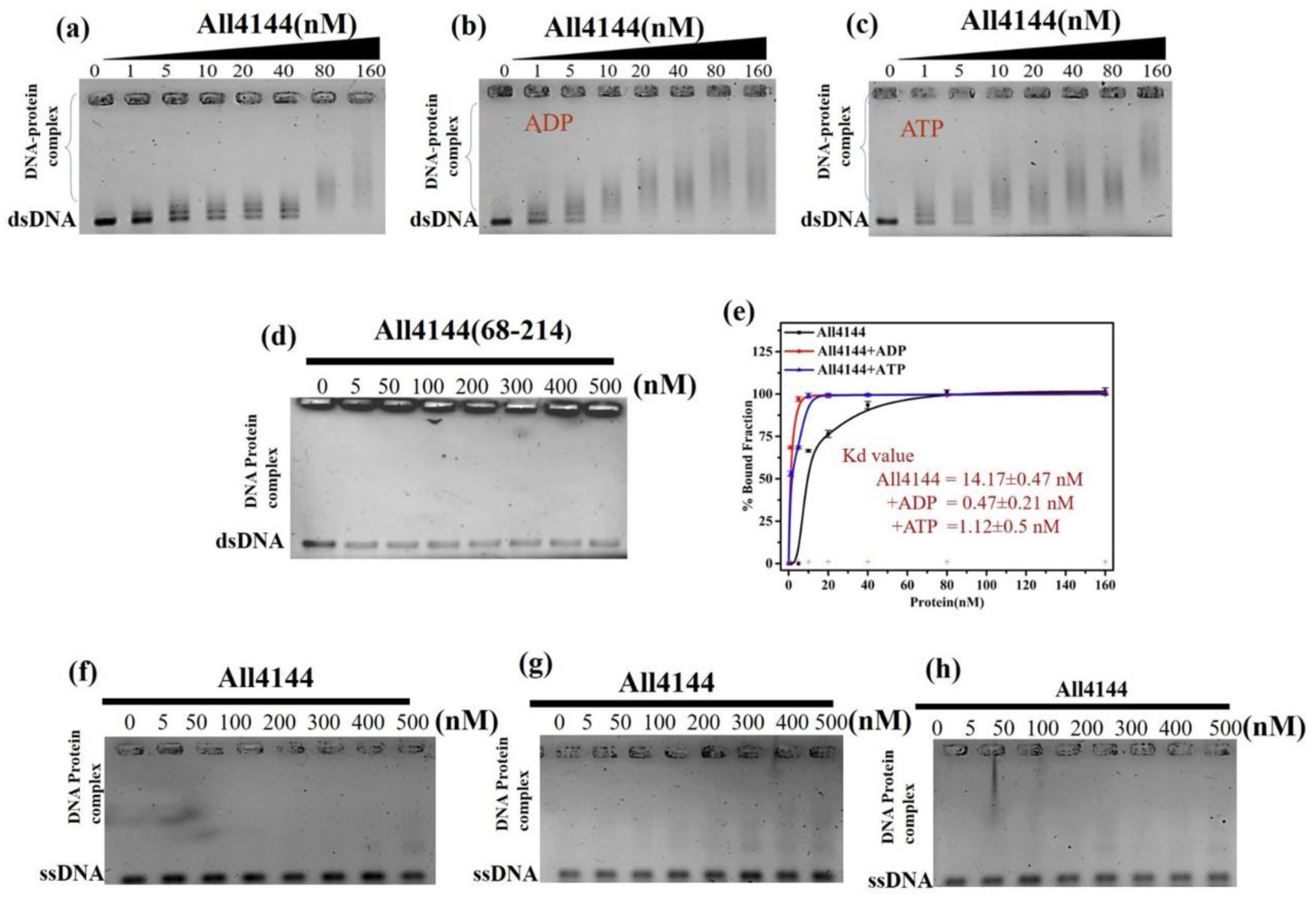
DNA-Protein Interaction Activity of All4144 and All4144(68–214): **(a)** dsDNA (501 bp) was subjected to incubation with varying concentrations (0, 1, 5, 10, 20, 40, 80, and 160 nM) of All4144 in the absence of ADP/ATP. **(b**) In the presence of 1 mM ADP and **(c)** 1 mM ATP, respectively **(d)** Similarly, increasing concentrations of All4144(68–214) were incubated with linear nonspecific dsDNA, titrated with different concentrations (0, 5, 50, 100, 200, 300, 400, and 500 nM) of All4144(68–214). **(e)** Densitometric estimation of band intensity for free and bound DNA, plotted as mean ± SD (n = 3) of the bound fraction as a percentage. The percentage of bound DNA for each concentration was calculated relative to the control (without protein). **(f-g)** For ss-DNA-protein interaction, increasing concentrations of All4144 were incubated with linear nonspecific ssDNA, titrated with 2-5 fold higher molar concentrations of All4144. Nucleoprotein complexes were electrophoresed in 1% agarose gel and stained with ethidium bromide. Data are presented as mean ± SD (n = 3). Statistical analysis using Student’s t-test was performed, and P values obtained at 95% confidence intervals are indicated as (*) for <0.001.

### All4144 has Unique Sequence Feature

Our study involved a comprehensive examination of sequence alignment, focusing on GrpE family proteins sourced from various organisms, including *E. coli*, *Arabidopsis*, human, rat, and several cyanobacterial strains. Our study encompassed a diverse range of cyanobacterial strains, such as *Roholtiella* sp., *Atlanticothrix silvestris*, *Nostoc punctiform*, *Trichormus variabilis*, *Fischerella thermalis*, *Calothrix* sp., and *Scytonema* sp. Additionally, we compared All4144 with its paralogs All4473 and Alr2445 from *Anabaena* sp., unveiling sequence identities of 14% and 18%, respectively. The proteins selected for multiple sequence alignment are detailed in Table 1, facilitating a comprehensive analysis of sequence conservation and divergence. Through a detailed examination of the alignment, we observed conserved features predominantly in the C-terminal region across all explored species, signified by a dark red star (box 2) in (Fig. 6a). Conversely, numerous notable sequence gaps were identified in the N-terminal region, highlighted by black stars (box 1) in (Fig. 6a). Our multiple sequence alignment (MSA) analysis highlights distinctive sequence features of All4144 compared to its paralogs (All4473 and Alr2445), providing valuable insights into the functional characteristics and evolutionary divergence within the GrpE family.

**Figure 6.**
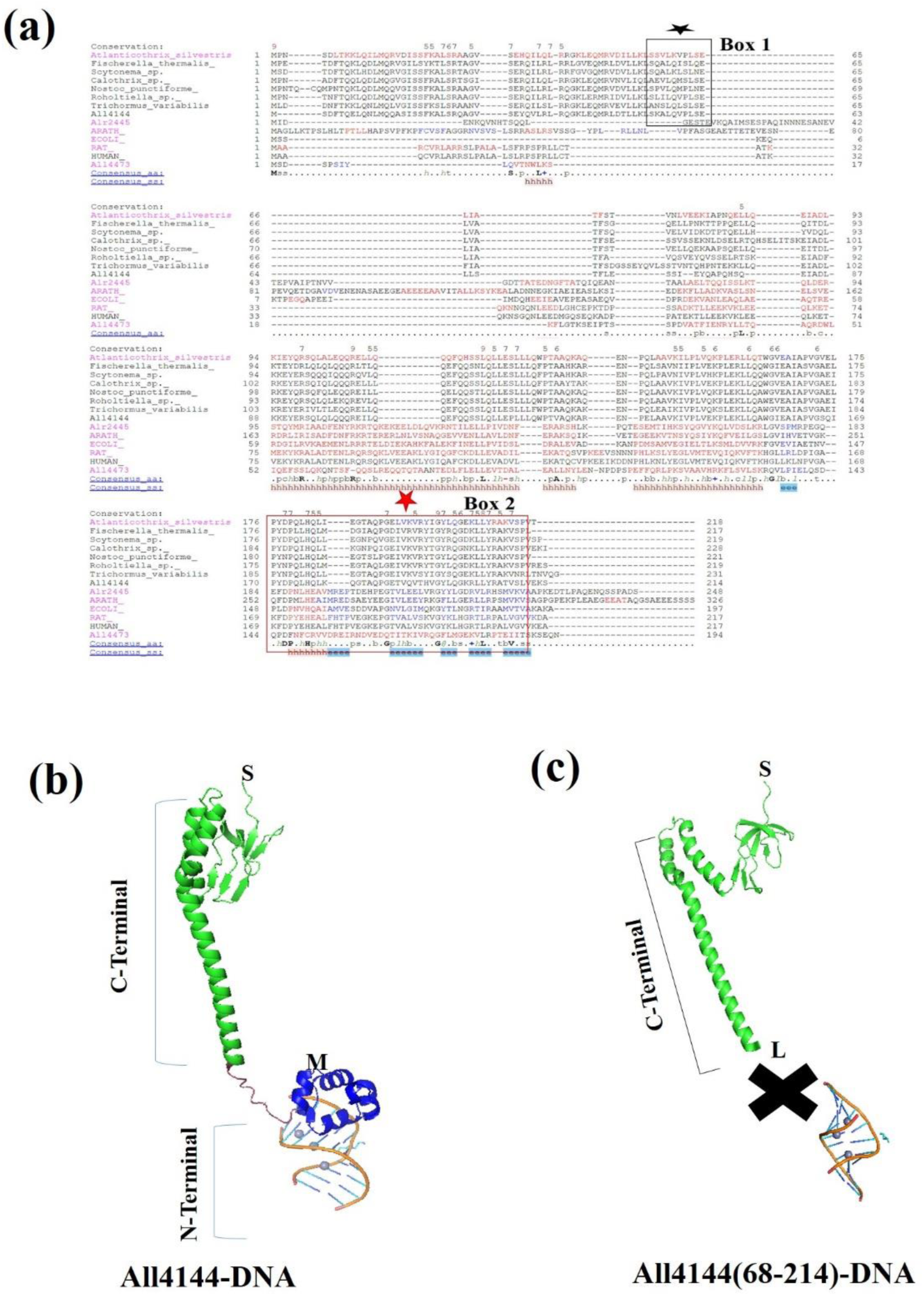
In-silico analysis of All4144, putative GrpE of Anabaena sp. PCC7120. **(a)** Sequence alignment of All4144 with other members of GrpE proteins from diverse organisms (accession number) using PROMALS-3D multiple sequence: N terminal region of All4144, marked by black star and C terminal region of All4144, marked by red star. **(b)** 3D model of All4144 interacts with DNA using HDOCK. **(c)** 3D model of All4144(68–214) and DNA.

**Table 1:**
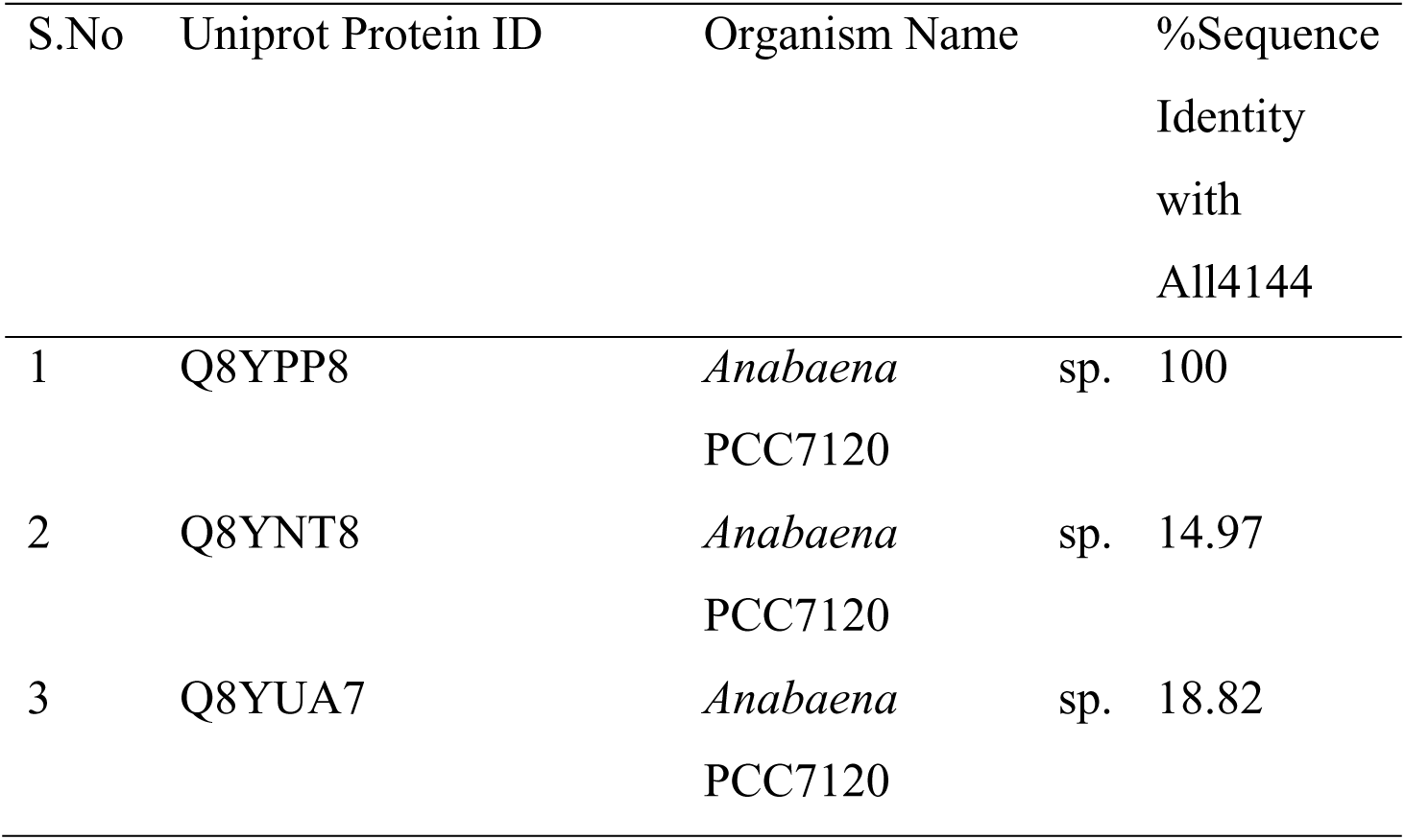
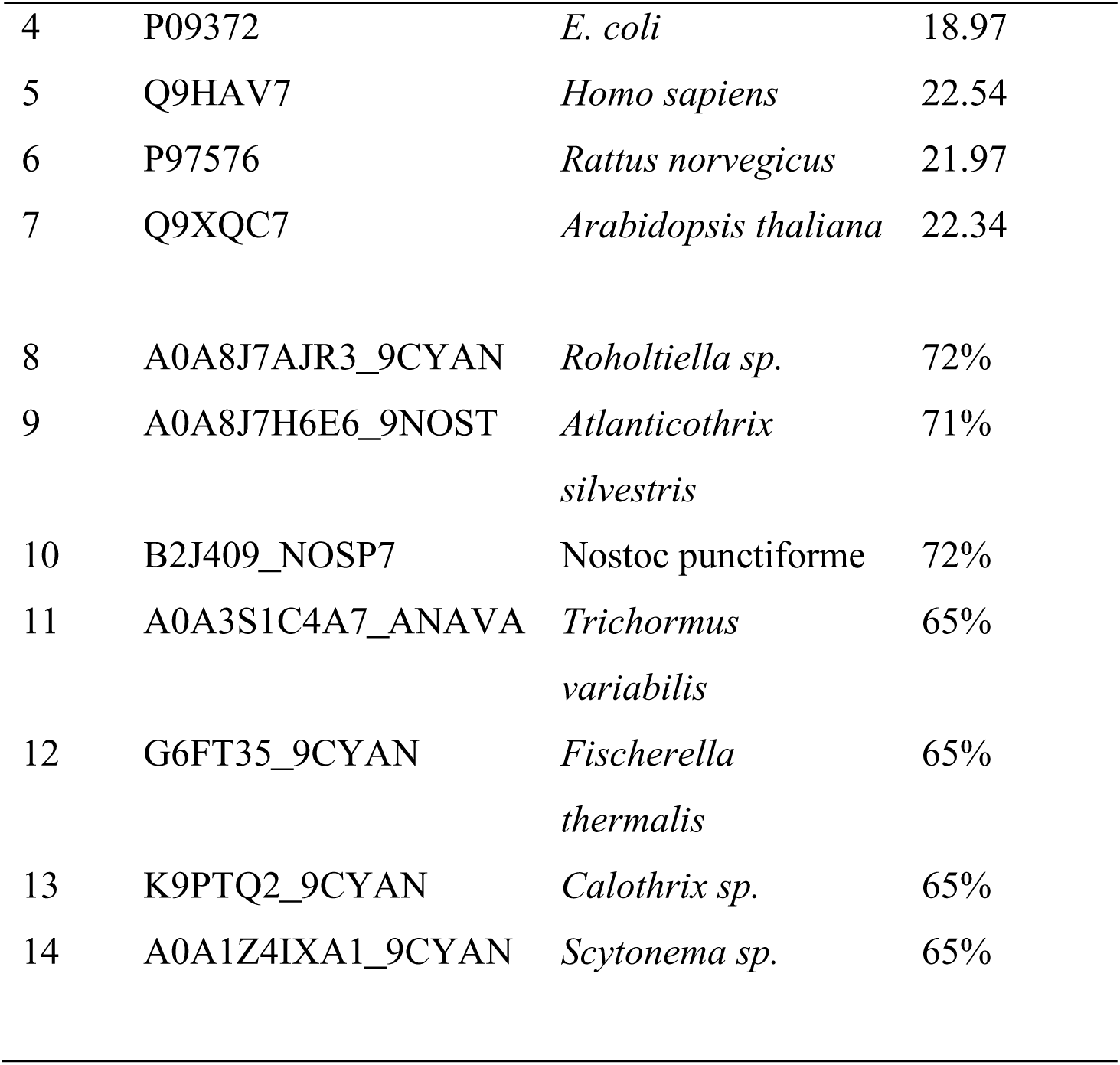
List of putative All4144 and GrpE family from other species, protein sequence ID and name used in multiple sequence alignMent.

| S.No | Uniprot Protein ID | Organism Name | %Sequence Identity with All4144 |
| --- | --- | --- | --- |
| 1 | Q8YPP8 | *Anabaena* sp. PCC7120 | 100 |
| 2 | Q8YNT8 | *Anabaena* sp. PCC7120 | 14.97 |
| 3 | Q8YUA7 | *Anabaena* sp. PCC7120 | 18.82 |
| 4 | P09372 | *E. coli* | 18.97 |
| 5 | Q9HAV7 | *Homo sapiens* | 22.54 |
| 6 | P97576 | *Rattus norvegicus* | 21.97 |
| 7 | Q9XQC7 | *Arabidopsis thaliana* | 22.34 |
| 8 | A0A8J7AJR3_9CYAN | *Roholtiella sp.* | 72% |
| 9 | A0A8J7H6E6_9NOST | *Atlanticothrix*<br>*silvestris* | 71% |
| 10 | B2J409_NOSP7 | *Nostoc punctiforme* | 72% |
| 11 | A0A3S1C4A7_ANAVA | *Trichormus*<br>*variabilis* | 65% |
| 12 | G6FT35_9CYAN | *Fischerella*<br>*thermalis* | 65% |
| 13 | K9PTQ2_9CYAN | *Calothrix sp.* | 65% |
| 14 | A0A1Z4IXA1_9CYAN | *Scytonema sp.* | 65% |

### Protein modeling and interaction with DNA

To elucidate the molecular characteristics of All4144, three-dimensional (3D) structure predictions of the full-length protein All4144 and its deletion mutant All4144(68–214) have been obtained from All4144 AlphaFold entry Q8YPP8. The structural integrity of the models was confirmed, and their quality was evaluated using RMSD (root-mean-square deviation), RAMPAGE and TM score. Subsequently, the HDOCK web server’s intuitive interface was employed to explore the interactions between the protein and DNA. The three-dimensional model of All4144 revealed that the N-terminal region of the protein is predicted to be involved in interactions with DNA (Fig. 6b). Notably, the deletion of the helix-turn-helix (HTH) domain from All4144 resulted in the loss of its ability to interact with DNA (Fig. 6c). This observation corroborates the results obtained from the electrophoretic mobility shift assay (EMSA), where the deletion of the N-terminal region completely abolished the protein’s DNA binding activity.

These findings collectively underscore the critical role of the N-terminal HTH domain in facilitating the interactions between All4144 and DNA. The integrated approach of structural modeling and experimental validation enhances our comprehension of the molecular mechanisms governing the DNA binding activity of All4144.

## 4. Discussion

The intricate regulation of chaperone systems across diverse organisms, encompassing the eukaryotic Hsc70/Bag1 and the prokaryotic DnaK/GrpE systems, has been extensively studied (38, 39, 40). The regulatory mechanisms within the DnaK chaperone system are complex, involving the interplay between ATP and ADP states, along with substrate release mediated by GrpE (41). GrpE plays a critical role in maintaining the nucleotide-binding domain (NBD) in the Hsp70/DnaK chaperone system, acting as an essential component for nucleotide exchange in DnaK (42).

Our previous findings (15) highlighted the ability of bacterial DnaJ-III to enhance ATP hydrolysis even in the absence of GrpE, prompting us to further investigate the influence of GrpE on the DnaK-DnaJ chaperone system and stress resilience. The current study focuses on All4144, an unconventional GrpE protein containing both HTH and GrpE domains, and its co-chaperone activity. We aim to understand how All4144 modulates protein folding and cellular stress responses. By exploring its influence on DnaK-DnaJ interactions, we seek novel insights into the regulatory mechanisms governing protein quality control and cellular homeostasis.

Our computational study revealed the presence of an N-terminal helix-turn-helix (HTH) domain in All4144, suggesting its potential role in DNA binding. This finding indicates that All4144 may possess additional functions beyond its conventional role as a co-chaperone for DnaK. Comparative sequence analysis highlighted significant conservation between All4144 and its mammalian counterparts. Additionally, previous findings (43) suggest that All4144, akin to Hap46, has dual functionalities: it binds DNA via its N terminus while acting as a nucleotide exchange factor for DnaK through its C terminus. This dual role implies that All4144 likely contributes to various metabolic processes in cyanobacteria. Moreover, the resemblance between All4144 and mammalian GrpE proteins indicates evolutionary convergence, hinting at similar functional roles across different organisms despite their evolutionary distance.

Transcript expression profiling revealed a significant increase in All4144 expression in response to heat stress, underscoring its role as a gene responsive to elevated temperatures. This suggests that All4144 is important for cellular responses to heat. Furthermore, cell survival and thermal assays demonstrated that All4144 significantly enhances heat stress tolerance.

GrpE’s function as a nucleotide exchange factor for DnaK is heat-inducible; it has been suggested that All4144 acts as a thermometer in response to elevated temperatures, as supported by previous findings (12, 44, 45).

Moreover, the dimerization properties of All4144, akin to other GrpE proteins further emphasize its role as a bona fide member of the GrpE family (46, 47). The observed structural features, coupled with the thermal transition profiles observed, suggest that All4144 and deletion mutant function as a true GrpE protein by modulating the kinetics of DnaK’s ADP/ATP conversion in response to temperature fluctuations. Additionally, the functional assays provided valuable insights into the interaction between All4144 and All4144(68–214) with DnaK, emphasizing its significance in modulating DnaK’s activity. This increased affinity likely enhances DnaK’s efficiency in ATP to ADP exchange, a crucial step in its chaperone cycle, and facilitates the refolding of thermally labile proteins such as malate dehydrogenase (MDH). Specifically, the deletion mutant alone acts as a complete GrpE protein, analogous to those found in other organisms like *E. coli* GrpE (46). However, All4144 exhibits a higher affinity for DnaK compared to its deletion mutant counterpart, possibly due to an additional domain HTH or alterations in the conformational properties of the full-length All4144 protein. This was confirmed by tryptophan fluorescence studies, which also suggested that the All4144 and DnaK-ADP complex more effectively quenched DnaK compared to its deletion mutant. This underscores the significance of both All4144 and its deletion mutant in the cellular stress response and chaperone machinery, contributing to the understanding of GrpE-mediated protein folding and stress adaptation mechanisms (48, 49).

It is noteworthy that despite the deletion of the N-terminal region, the interaction between All4144 and DnaK remained intact, and its nucleotide exchange factor activity was preserved. This observation is consistent with previous findings regarding other GrpE proteins, suggesting that the N-terminal region may not be essential for the GrpE-DnaK interaction or its nucleotide exchange factor activity (50).

Furthermore, the predictions regarding protein-DNA interactions, along with the results of electrophoretic mobility shift assays (EMSAs), strongly indicate the involvement of the N-terminal region of All4144 in DNA interaction. This observation suggests a potentially novel role for All4144 protein in mediating DNA interactions, particularly through their N-terminal domains. The preference of All4144 for binding to double-stranded (dsDNA) and not bind with ssDNA, especially higher in the presence of ADP, further underscores the possibility of GrpE acting as a nucleotide exchange factor in DNA interactions. This implies a multifaceted role for All4144 in cellular processes beyond conventional chaperone functions, expanding our understanding of the diverse functions of GrpE proteins in cellular physiology and stress response pathways. Further exploration of these interactions will provide valuable insights into the intricate molecular mechanisms underlying DNA-protein interactions mediated by GrpE proteins.

## Conclusion

All4144 emerges as a novel GrpE protein with distinct characteristics, deviating from its counterparts. The inclusion of the HTH domain enhances its functionality and augments thermal tolerance, shedding light on its multifaceted roles in modulating the DnaK chaperone cycle and interacting with DNA. Future investigations will elucidate the specific contributions of these domains to cyanobacterial physiology.

## Acknowledgments

L.C. Rai thanks National Academy of Science, India for the NASI Senior Scientist Platinum Jubilee Fellowship and the Indian Council of Agricultural Research-National Bureau of Agriculturally Important Microorganisms (ICAR-NBAIM) for financial support. Sonam Sriwastaw is thankful to the Department of Science and Technology (DST) for the Senior Research Fellowship (SRF).

## CRediT authorship contribution statement

SS and LCR designed the experiments. SS conducted the experiment. SS and RR have written the manuscript. SS and LCR analyzed the data. LCR critically reviewed the paper.

## References

1. A.K. Singh, P.P. Singh, V. Tripathi, H. Verma, S.K. Singh, A.K. Srivastava, A. Kumar. 2018. Distribution of cyanobacteria and their interactions with pesticides in paddy field: a comprehensive review *J*. Environ. Manag., 224, pp. 361–375.

2. Babele, P.K., Kumar, J., Chaturvedi, V. 2019. Proteomic De-regulation in cyanobacteria in response to abiotic stresses. Front. Microbiol. 10, 1–22.

3. Tyedmers, J., Mogk, A., Bukau, B. 2010. Cellular strategies for controlling protein aggregation. Nat. Rev. Mol. Cell Biol. 11, 777–788.

4. Ghazaei C. 2017. Role and mechanism of the Hsp70 molecular chaperone machines in bacterial pathogens. Microbiol Res 259–265.

5. McDonald HJ, Kweon HJ, Kurnfuli S, Risser DD. 2022. A DnaK(Hsp70) chaperone system connects type IV pilus activity to polysaccharide secretion in cyanobacteria. MBio 13: e0051422

6. Kampinga HH. 2019 The HSP70 chaperone machinery: J proteins as drivers of functional specificity. Nat Rev Mol Cell Biol. 11(8): 579–592.

7. Zuiderweg ERP, Hightower LE, Gestwicki JE. 2017. The remarkable multivalency of the Hsp70 chaperones. Cell Stress Chaperones 22:173–189.

8. Liu J, Chen L, Wang J et al. 2012. Proteomic analysis reveals resistance mechanism against biofuel hexane in *Synechocystis* sp. PCC 6803. Biotechnol. Biofuels 5:68.

9. Marada A, Karri S, Singh S et al. 2016 A single point mutation in mitochondrial Hsp70 cochaperone Mge1 gains thermal stability and resistance. Biochemistry 55:(51),7065–7072.

10. Naylor DJ, Stines AP, Hoogenraad NJ, Høj PB. 1998. Evidence for the existence of distinct mammalian cytosolic, microsomal, and two mitochondrial GrpE-like proteins, the co-chaperones of specific Hsp70 members. J Biol Chem 273:21169–21177.

11. Sondermann H, Ho AK, Listenberger LL, et al. 2002. Prediction of novel Bag-1 homologs based on structure / function analysis identifies Snl1p as an Hsp70 co-chaperone in *Saccharomyces cerevisiae*. J Biol Chem 277:33220–33227.

12. Barthel S, Rupprecht E, Schneider D. 2011. Thermostability of two cyanobacterial GrpE thermosensors. Plant Cell Physiol 52:1776–1785.

13. Gelinas AD, Toth J, Bethoney KA, et al. 2004. Mutational analysis of the energetics of the GrpE · DnaK binding Interface : Equilibrium association constants by sedimentation velocity analytical ultracentrifugation. Boston Biomedical Research 447–458.

14. Nakamura A, Takumi K, Miki K. 2010. Crystal structure of a thermophilic GrpE protein : Insight into thermosensing function for the DnaK chaperone system. J Mol Biol 396:1000–1011.

15. Sriwastaw S, Rai R, Raj A, et al. 2023. All3048, a DnaJ III homolog of *Anabaena* sp. PCC7120 mediates heat shock response in *E. coli* and its N-terminus J-domain stimulates DnaK ATPase activity. Int J Biol Macromol 123563.

16. Sambrook J. 2001. Molecular cloning : a laboratory manual, 3rd ed. Cold spring harbor laboratory press, *Cold Spring Harbor* N.Y. 56

17. Quevillon E, Silventoinen V, Pillai S, et al. 2005. InterProScan: Protein domains identifier. Nucleic Acids Res 33:116–120.

18. Marchler-Bauer A, Lu S, Anderson JB, et al. 2011. CDD: A conserved domain database for the functional annotation of proteins. Nucleic Acids Res 39:225– 229.

19. Kumar S, Stecher G, Li M, et al. 2018. MEGA X: Molecular evolutionary genetics analysis across computing platforms. Mol Biol Evol 35:1547–1549.

20. Yang J, Yan R, Roy A, et al. 2015. The I-TASSER Suite: Protein structure and function prediction. Nature Methods 12:7–8.

21. Ramesh DMK V. 2020. Binding site analysis of potential protease inhibitors of COVID-19 using AutoDock. VirusDisease 31:194–199.

22. Yan, Y., Tao, H., He, J., & Huang, S. Y. (2020). The HDOCK server for integrated protein–protein docking. Nature protocols, 15(5), 1829–1852.

23. Pandey, S., Rai, R., Rai, L.C., 2012. Proteomics combines morphological, physiological and biochemical attributes to unravel the survival strategy of Anabaena sp. PCC7120 under arsenic stress. J. Proteomics 75, 921–937.

24. Rajaram, H., Apte, S.K. 2008. Nitrogen status and heat-stress-dependent differential expression of the cpn60 chaperonin gene influences thermotolerance in the cyanobacterium Anabaena. Microbiology 154, 317– 325.

25. Rai, S., Agrawal, C., Shrivastava, A.K., Singh, P.K., Rai, L.C. 2014. Comparative proteomics unveils cross species variations in Anabaena under salt stress. J. Proteomics 98, 254–270.

26. Rai, S., Singh, S., Shrivastava, A.K., Rai, L.C. 2013. Salt and UV-B induced changes in *Anabaena* PCC 7120: Physiological, proteomic and bioinformatic perspectives. Photosynth. Res. 118, 105–114.

27. Singh, P.K., Shrivastava, A.K., Chatterjee, A., Pandey, S., Rai, S., Singh, S., Rai, L.C. 2015 Cadmium toxicity in diazotrophic *Anabaena* sp. adjudged by hasty up-accumulation of transporter and signaling and severe down-accumulation of nitrogen metabolism proteins. J. Proteomics 127, 134–146.

28. Livak, K.J., Schmittgen, T.D. 2001. Analysis of Relative Gene Expression Data Using Real-Time Quantitative PCR and the 2−ΔΔCT Method. Methods 25, 402–408.

29. Agrawal C, Sen S, Yadav S, et al. 2015. A novel aldo-keto reductase (AKR17A1) of *Anabaena* sp. PCC 7120 degrades the rice field herbicide butachlor and confers tolerance to abiotic stresses in *E. coli*. PLOS ONE 10:9

30. Jakubowski W, Bartosz G. 2000. 2’,7’-Dichlorodihydro fluorescein as a fluorescent probe for reactive oxygen species measurement: Forty years of application and controversy. Cell Biol Int. 24:757–760.

31. Wen J, Batabyal D, Knutson N, et al. 2020. A comparison between emerging and current biophysical methods for the assessment of higher-order structure of Biopharmaceuticals. J Pharm Sci 109:247–253.

32. Seelig, J.; Seelig, A. 2023. Protein Unfolding─Thermodynamic Perspectives and Unfolding Models. Int. J. Mol. 24, 5457.

33. Maurya GK, Kota S, Misra HS. 2019. Characterisation of ParB encoded on multipartite genome in *Deinococcus radiodurans* and their roles in radioresistance. Microbiol Res 223–225:22–32.

34. Veinger L, Diamant S, Buchner J, Goloubinoff P. 1998. The small heat-shock protein IbpB from *Escherichia coli* stabilizes stress-denatured proteins for subsequent refolding by a multichaperone network. J Biol Chem 273:11032– 11037.

35. P.A. Lanzetta, L.J. Alvarez, P.S. Reinach, O.A. Candia. 1979. An improved assay for nanomole amounts of inorganic phosphate, Anal. Biochem. 100 95–97,

36. Han W, Christen P. 2003. Interdomain communication in the molecular chaperone DnaK. Biochem J 369:627–634.

37. Wei X, Mingjia H, Xiufeng L, et al. 2007. Identification and biochemical properties of Dps (starvation-induced DNA binding protein) from cyanobacterium *Anabaena* sp. PCC 7120. IUBMB Life. 59:675–681.

38. Johnston, C.L., Marzano, N.R., van Oijen, A.M., Ecroyd, H. 2018. Using Single-Molecule Approaches to Understand the Molecular Mechanisms of Heat-Shock Protein Chaperone Function. J. Mol. Biol. 430, 4525–4546.

39. T.J. Lupoli, A. Fay, C. Adura, M.S. Glickman, C.F. Nathan. 2016. Reconstitution of a Mycobacterium tuberculosis proteostasis network highlights essential cofactor interactions with chaperone DnaK . Proc. Natl Acad. Sci. USA, 113 (49), pp. E7947–E7956.

40. Srivastava, S., Savanur, M.A., Sinha, D., Birje, A., Vigneshwaran, R., Saha, P.P., Silva, P.D. 2017. cro Regulation of mitochondrial protein import by the nucleotide exchange factors GrpEL1 and GrpEL2 in human cells J Biol Chem 292, 18075–18090.

41. Preissler S, Chambers JE, Crespillo-casado A, et al. 2015. Physiological modulation of BiP activity by trans -protomer engagement of the interdomain linker. elifesciences 1–31.

42. Imamoglu R, Balchin D, Hayer-Hartl M, Hartl FU. 2020. Bacterial Hsp70 resolves misfolded states and accelerates productive folding of a multi-domain protein. Nat Commun 11:365.

43. Niyaz Y, Frenz I, Petersen G, Gehring U. 2003. Transcriptional stimulation by the DNA binding molecular chaperones. Nucleic Acids Res 31:2–9.

44. A.D. Gelinas, J. Toth, K.A. Bethoney, K. Langsetmo, W.F. Stafford, C.J. Harrison 2003. ,Thermodynamic Linkage in the GrpE Nucleotide Exchange Factor, a Molecular Thermosensor. Biochemistry, 42 pp. 9050–9059.

45. Upadhyay, T., Karekar, V. V., Potteth, U. S., & Saraogi, I. 2023 Investigating the functional role of a buried interchain aromatic cluster in Escherichia coli GrpE dimer. Proteins: Structure, Function, and Bioinformatics, 91(1), 108–120.

46. Harrison C. 2003 GrpE, a nucleotide exchange factor for DnaK. Cell Stress Chaperones 8 (14984054): 218–224.

47. Moro, F., Taneva, S.G., Velázquez-campoy, A., Muga, A. 2007. GrpE N-terminal Domain Contributes to the Interaction with DnaK and Modulates the Dynamics of the Chaperone Substrate Binding Domain. J. Mol. Biol 7;374(4): 1054–1064.

48. L. Rohland, R. Kityk, L. Smalinskaitė, M.P. Mayer. 2022. Conformational dynamics of the Hsp70 chaperone throughout key steps of its ATPase cycle. Proc Natl Acad Sci U S A, 119 : e2123238119

49. Xu H. 2018. Cochaperones enable Hsp70 to use ATP energy to stabilize native proteins out of the folding equilibrium. Sci Rep 8:1–15.

50. Liberek, K., Marszalek, J., Ang, D., Georgopoulos, C. & Zylicz, M. 1991. Escherichia coli DnaJ and GrpE heat shock proteins jointly stimulate ATPase activity of DnaK. Proc. Natl Acad. Sci. USA, 88, 2874–2878.

